# Dataset Augmentation Allows Deep Learning-Based Virtual Screening To Better Generalize To Unseen Target Classes, And Highlight Important Binding Interactions

**DOI:** 10.1101/2020.03.06.979625

**Authors:** Jack Scantlebury, Nathan Brown, Frank Von Delft, Charlotte M. Deane

**Affiliations:** Department of Statistics, University of Oxford, 24-29 St Giles, Oxford, OX1 3LB, UK; BenevolentAI, 4-8 Maple St, London, W1T 5HD, UK; Structural Genomics Consortium (SGC), University of Oxford, Oxford, OX3 7DQ, UK; Diamond Light Source, Harwell Science and Innovation Campus, Didcot OX11 0DE, UK; Department of Biochemistry, University of Johannesburg, Aukland Park, Johannesburg 2006, South Africa

## Abstract

Current deep learning methods for structure-based virtual screening take the structures of both the protein and the ligand as input but make little or no use of the protein structure when predicting ligand binding. Here we show how a relatively simple method of dataset augmentation forces such deep learning methods to take into account information from the protein. Models trained in this way are more generalisable (make better predictions on protein-ligand complexes from a different distribution to the training data). They also assign more meaningful importance to the protein and ligand atoms involved in binding. Overall, our results show that dataset augmentation can help deep learning based virtual screening to learn physical interactions rather than dataset biases.

**Graphical TOC Entry:** 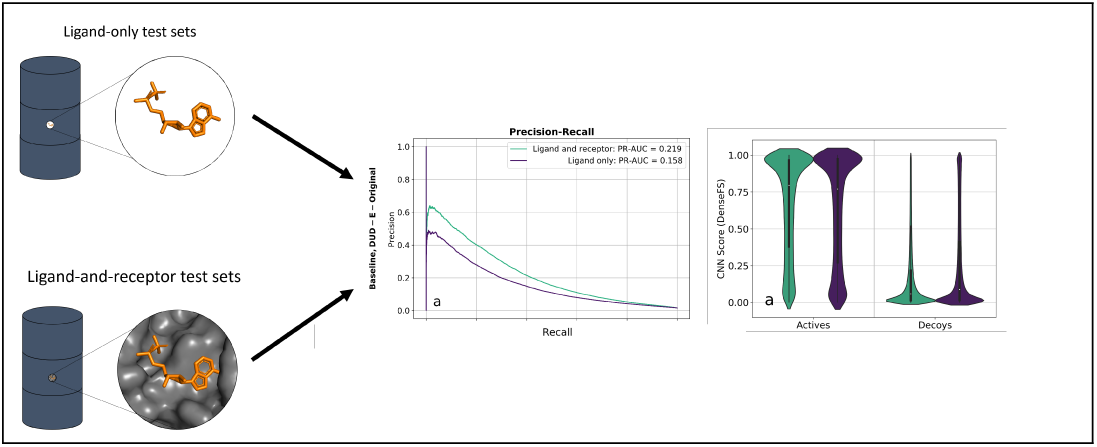

## Introduction

A series of recent papers has shown that some deep learning methods designed for structure-based virtual screening can accurately separate actives and decoys when given only the structure of the ligand.^1–3^ These results indicate that such methods are learning differences between the properties of actives and decoys, rather than the physical interactions between the receptor and the ligand. From this it is possible to conclude both that the methods will fail to generalize well (predict on datasets far removed from the training data), and that there are significant flaws in the current training datasets and/or regimens.

The task of a virtual screening algorithm is to distinguish between bound structures where the small molecule is an active, and those where it is a decoy. A standard structure-based virtual screening technique docks potential drug-like compounds into a protein target of interest, and ranks them according to a physically-inspired scoring function (for a review of common virtual screening techniques including docking, see [4]). These methods can be used to screen vast numbers of potential compounds for target interactions and are extremely cheap compared to lab-based experiments. ^5^ The accuracy of docking, however, is highly target-dependent.^6^ This issue, along with the current deluge of structural data, has prompted the use of machine learning methods in structure-based virtual screening and in the verification of docked structures. ^7–9^

Machine learning has found its way into many scientific domains, and drug discovery is no exception (a review can be found in [10]). Multiple machine learning-based virtual screening methods have been developed. Most of these rely on 1- or 2-D descriptors - that is, they take as input a representation of the ligand as a fingerprint,^11^ graph^12,13^ or other descriptors, ^14^ and the protein (if present at all) as a sequence of amino acids.^13^ More recently, machine learning methods which are able to capture specific spatial and chemical interactions between the protein and ligand have been introduced; the major driver for building these more complex methods is the hope that because they can learn physicochemical interactions between protein and ligand, they should be able to more accurately predict the binding of ligands and proteins far removed from the complexes on which they were trained (be more generalisable). The current state-of-the-art method for attempting to capture these types of spatial interactions is the convolutional neural network (CNN).^15^

CNNs are a type of deep neural network which, due to their ability to capture spatial relationships between objects, are commonly used in image classification. The gnina framework for virtual screening^16^ treats docked structures as 3D images with different channels for different types of atoms as inputs into a CNN and outputs a binary classification (active/decoy). A CNN with three hidden layers built in the gnina framework and trained and tested on part of the DUD-E dataset^17^ was shown to perform better in the task of virtual screening than using the docking score of AutoDock Vina. It had a mean test target ROC-AUC of 0.862, compared to 0.703 for AutoDock Vina (reported in [18]); a ROC-AUC score is the area under the curve of the receiver-operator characteristic graph, which is a measure of the ability of a classifier to correctly rank unseen labelled data. The same input format with a deeper and more densely connected network along with protein family-specific finetuning was used to develop the DenseFS CNN, which significantly increased predictive power to a mean DUD-E target ROC-AUC of 0.917.^18^ This featurization method has also been used to explore the nature and role of hydration in protein-ligand binding.^19^

These results suggest that deep learning offers real potential in terms of structure-based virtual screening. However, both CNN methods performed far worse on validation sets taken from a different database of structures (they generalize poorly). For example in the case of DenseFS, models trained on the DUD-E dataset and tested on an external dataset constructed from the ChEMBL database^20^ performed significantly worse by every metric (mean test target ROC-AUC and mean precision) than when the same models were tested on DUD-E.^18^

It has been demonstrated many times that a machine learning model with good performance on held-out test data from the same distribution as the training data does not guarantee good performance on other data sets. ^1,2,18^ This lack of generalisability means that models cannot accurately classify ligands which are significantly larger or smaller than the molecules they were trained on, or ligands which are chemically distinct from the training data.

If the aim of generalisability to targets far from the training set is to be realized, it is important that structure-based virtual screening learn physical interactions between the protein and the ligand, rather than properties of the ligand. One way to identify if methods are learning such interactions is to visualize the importance of atoms or groups of atoms to the score. Hochuli *et al.* used several techniques to visualize which atoms and residues in the input to the original gnina network were important to the classification decision. ^21^ One method used was input masking, where the CNN is used to score the docked complex both with and without certain atoms or groups of atoms present. These scores can then be compared in order to ascertain the importance of these atoms or groups of atoms.

There have also been attempts to quantify what precisely is being learned by CNNs for virtual screening. It was recently shown that model performance for some methods does not suffer significantly when ligand structures in the test set are given without the protein target/receptor.^2,3^ The overall conclusion from these studies was that the methods are using very little if any information from the protein target. This finding links to the fact that in many of these datatsets, receptor-free methods such as k-nearest neighbours on ligand fingerprints perform almost as well as methods making use of the receptor.^1^ It has even been shown that using a small number of simple 1D properties of ligand molecules is enough to classify accurately on the widely-used DUD-E dataset. The 50 decoys per active in the DUD-E dataset were chosen to have the same or similar values as the active molecule for six key properties (molecular weight, ClogP, number of hydrogen bond acceptors and donors, net charge and number of rotatable bonds) whilst remaining topologically distinct; classifiers trained using just these six supposedly unbiased properties achieved good performance on held-out DUD-E targets.^2^ This result indicates that differences between the properties of actives and decoys in the DUD-E dataset without any protein information can be used for classification.

The high performance of CNNs for virtual screening when only given the ligand structure and the known biases in the dataset suggests that networks are not learning the physico-chemical interactions between the protein and the ligand, but are separating actives from decoys based on properties of the ligand alone.

A recent attempt to remedy this problem was made using both Maxiumum Unbiased Validation (MUV) and Asymmetric Validation Embedding (AVE). MUV is a measure of how clustered actives are and AVE measures how tightly clustered decoys are relative to the active-decoy distance. In [22], the distance between molecules was defined using their 2048-bit ECFP6 fingerprint, and ‘unbiased’ training sets were generated by minimising either MUV or AVE with a genetic algorithm. The training data was 189 targets each with at least 500 actives. Two types of tests sets were generated: ‘standard-AUC’ sets consisting of randomly selected (non-training) ligands, and ‘far-AUC’ sets which consist of ligands that are considered to be far (Jaccard dissimilarity between ECFP6 fingerprints ≥ 0.4) from ligands in the training set for that target. It was found that the original (‘biased’) models which performed well on standard-AUC often failed to perform as well on far-AUC, showing the expected lack of generalisability. However, training on the unbiased training sets led to a drop in performance on far-AUC, meaning that this type of debiasing is ineffective for improving generalisability.

In this paper, we test how CNNs use receptor information and explore methods to improve their generalisability. First, we compare the behaviour of deeper CNNs to their shallower counterparts, and find that deeper CNNs use more information from the receptor, but their classifications are still heavily reliant on the identity of the ligand. Given this result, we develop a procedure to force the CNN to learn from the protein/ligand interactions. We augment the training dataset with active ligands incorrectly positioned and labelled as decoys. In order to analyse the effect of this augmentation, we look both at the ability of the method to predict and the distribution of CNN scores attained for different training conditions. On cross-validation (training and testing on different parts of the DUD-E dataset), there is no significant difference in performance between the original and new methods, but the score distributions show increased sensitivity to receptor information when training data is augmented in this way. Finally, to demonstrate the method trained with this augmented dataset is learning physical interactions, we show both that the method is more generalisable, it performs significantly better on an external test set, and that atoms involved in interactions between the ligand and receptor are now important to the score.

## Methods

To probe what is being learned by CNNs for virtual screening and to develop methods to force the learning of physical interactions, three facets of the problem were explored. These were: the network architecture (model), the training set, and the presence or otherwise of receptor information in the test sets.

### Datasets

All training was carried out using the Database of Useful Decoys - Extended (DUD-E). ^17^ To carry out initial tests, a three-fold cross validation strategy on DUD-E was used. External validation was carried out using the same subset of targets and ligands from the ChEMBL dataset^20^ as in the previous study of Imrie *et al.*^18^

### DUD-E

The Database of Useful Decoys Extended (DUD-E) ^17^ is comprised of more than 22,000 actives and 1,100,000 decoys across 102 protein targets. Actives are molecules which have been experimentally determined to bind in the binding pocket of a protein target, and for each active, there are 50 decoys which are assumed to be non-binders. All actives and decoys are docked into the binding site using AutoDock Vina. ^23^ The decoys in DUD-E were generated to have similar physicochemical properties to their active (molecular weight, log-P, number of hydrogen bond acceptors and donors, net charge and number of rotatable bonds), but were designed to be topologically distinct.

Following the methodology of Imrie *et al.*,^18^ three folds were constructed from DUD-E where protein targets with ≥ 80% sequence identity were placed in the same fold, to avoid training on structures that are similar to those in the test fold. The details of these folds can be found in the supporting information (Table S1). Intra-dataset validation was carried out by training on two folds with the remaining one used for testing, giving three possible permutations.

Each DUD-E target can also be categorized into one of the following families, with numbers indicating the number of targets per category: Kinase (26), Protease (15), Nuclear (11), G protein-coupled receptor (GPCR) (15) and Other (45).

### ChEMBL

A set of 50 ChEMBL targets determined to be a suitable benchmark set for 2D fingerprinting methods was compiled by Heikamp and Bajorath. ^24^ Of these, a subset of 14 targets which had ≤ 80% sequence similarity to all DUD-E targets were chosen as an external validation set in both Ragoza *et al.*^16^ and Imrie *et al.*.^18^ There are nine docked structures for each active, all of which are labelled as active. For each active ligand, there are *∼*100 decoy ligands, again each with nine docked structures labelled as decoys. This gives 12,275 active structures over 14 targets, with 1,363 unique active ligand molecules, along with 1,238,178 decoy structures.

### CNN Architectures

Gnina^16^ defines the binding site as a cube with sides of 24 °A centred on the ligand, which it splits into 48 × 48 × 48 voxels, each with sides of length 0.5 °A. It also constructs 34 channels, each containing the pseudo-electron density from a different atom type, and designated as either a ‘ligand channel’ (18) or ‘protein channel’ (16). The density at each voxel is calculated using a piecewise combination of a Gaussian and a quadratic function, both depending on the distance from the nucleus and the van der Waals radius. The combination of all 34 channels over the three dimensions gives a 48 × 48 × 48 × 34 tensor description of the input space which is used as the feature vector for the CNN. The two models described below both use this featurization method, but differ in the network architecture used for classification. The authors of Gnina have released a standalone version of *libmolgrid* ^25^ (the featurization part of Gnina), which can be found at https://gnina.github.io/libmolgrid.

#### Gnina

A network with three convolutional layers (each followed by a max pool), then a fully-connected final layer with a softmax activation function, giving a probability over the two categories (active/decoy). This version of the CNN ^16^ was the subject of all the analyses so far carried out into the importance of the receptor in active/decoy classification. ^1–3^

#### DenseFS

A much deeper network with three sets of four densely connected convolutional layers followed by a fully-connected softmax layer and cross entropy loss. ^18^ This network significantly improved performance over the Gnina network on both held-out DUD-E targets and the ChEMBL set.

### Training Schemes

Training was carried out using the original DUD-E dataset as well as three different augmented datasets:

- **DUD-E-Original** The original DUD-E dataset, using the docked poses provided in Ragoza *et al*.^16^
- **DUD-E-Trans** DUD-E-Original, augmented with three copies of each active, in a random conformation and translation. See Protocol S1 and Fig. S1 in the supporting information for the full protocol and a visual example.
- **DUD-E-Redocked** DUD-E-Original, augmented by taking each active, and redocking it into the binding pocket with AutoDock Vina to generate up to 20 poses. The three highest ranked (lowest energy) poses at least 5°A root-mean-squared distance (RMSD) from the active pose were labelled as decoys and used in training.
- **DUD-E-Hybrid** Both augmentations from DUD-E-Trans and DUD-E-Redocked were added to the training set, giving a total of six extra decoys per active.

### Models

From hereon in, **Baseline** will refer to models with the Gnina architecture trained on all or part of DUD-E-Original and **OriginalFS, TransFS, RedockedFS** and **HybridFS** to classifiers with the DenseFS architecture trained on all or part of DUD-E-Original, DUD-E-Trans, DUD-E-Redocked and DUD-E-Hybrid respectively.

### Ligand-and-Receptor and Ligand-Only Tests

The two tests described in this section (Ligand-and-receptor and Ligand-only) were conducted according to the same protocol, the only difference being the presence or absence of the receptor in the test set structures. As described above and following Imrie *et al.*,^18^ three folds were constructed from the 102 DUD-E targets (Table S1 in the supporting information). A model was trained on two of the three folds, with the final one reserved for testing. Test predictions from all three models were concatenated for easier visualisation of results. This protocol is depicted in Fig. S2 in the supporting information.

#### Models for the Ligand-and-receptor and Ligand-only tests

Three separate CNNs (one for each fold) were trained using each of the five CNN models described above, giving a total of 15 trained models. Training was conducted with a batch size of 16, for 25,000 iterations and the number of actives and decoys in each batch was even.^18^ The optimizer used was stochastic gradient descent, with a base learning rate is 0.01, an inverse learning rate power decay value of 1, a gamma value of 0.001, a weight decay of 0.001, with a momentum of 0.9.

#### Ligand-and-receptor Test

Once the models were trained according to the above protocol, they were tested on each of the test folds with no changes. This was the Ligand-and-receptor test.

#### The Ligand-only Test

A receptor-free version of each test fold was generated. For every case (actives and decoys) the protein was removed from the structure file leaving only the docked ligand in the pose it has in the protein binding pocket. This was the Ligand-only test.

### ChEMBL Validation

In order to investigate the effect of our training data augmentation on how CNNs generalize to data from a different distribution to the training set, three CNNs were trained on the entire DUD-E dataset (or an augmented version) with different random seeds. These CNNs were then used to classify actives and decoys in our ChEMBL validation set. Following Imrie *et al.*,^18^ the score assigned to each structure was the mean of the three CNN scores; the score assigned to a ligand with a target was the mean of the scores given to the top five (of the nine) structures. This system is illustrated in the supporting information (Fig. S3). All training was conducted with the same hyperparameters as outlined above.

In Imrie *et al.*,^18^ models were first trained on DUD-E in the same way as described in the methods section. However, the authors also made use of ‘finetuning’, a process of further tweaking all or part of the network for specific tasks. In this instance, copies of the initially trained networks were trained on different families on DUD-E, and then the test data was filtered according to family and the network such that the network that would be used to classify would be that which was finetuned on the training data from that family. This was reported to greatly improve discriminative power. We did not to employ that method, as the aim is only to test if augmentation improves generalisability of the model.

### Masking

Classifications from neural networks are notoriously difficult to interrogate. The masking method of Hochuli *et al.* attempts to obtain the contributions from different residues in the binding site, and different atoms of the ligand. ^21^ In short, a trained model is given a bound structure, and then the same structure with different ligand atoms or receptor residues removed (the structure is ‘masked’). The difference in CNN score between the full structure and the masked structure is taken as the contribution of whichever entity was removed (see Fig. S5 in the supporting information). This difference is then converted to a percentage of the original CNN score. The sum of all of the individual atomic contributions need not in general add to 100%, because the relationship between the input features and the CNN score is nonlinear.

In the original paper (and accompanying software), only residue contributions from the receptor were calculated. Here we modified the software to mask individual atoms of the protein, in order to gain a higher resolution image. The PDB^26^ structures used for masking (10OH and 1W4O) and were chosen to be the same as those used for masking by Hochuli *et al.* in [21] for ease of comparison. Neither of the PDB structures contains protein/ligand combinations found in DUD-E.

### Methods of Assessment

Two metrics were used to measure the discriminative power of the classifiers, the area under the curve for receiver-operator characteristic graph (ROC-AUC) and the area under the curve for the precision-recall curve (PR-AUC). While the ROC-AUC usually lies between 0.5 and 1.0 (for random and perfect classifiers), the PR-AUC will usually lie between 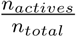 and 1.0, where *n*_*actives*_ is the number of actives in the test set and *n*_*total*_ is the total test set size. These lower bounds (which represent the performance of a random classifier) are *∼* 0.02 and *∼* 0.01 for the DUD-E test sets and ChEMBL validation sets respectively.

The use of ROC-AUC as a metric for model performance on highly imbalanced datasets such as DUD-E (50:1 actives:decoys) or the ChEMBL set used here (100:1) has been shown to be less useful than PR-AUC.^27^ ROC-AUC is widely reported for classification algorithms and is included here for ease of comparison with other work.

## Results and Discussion

Only results using models trained on DUD-E-Original (OriginalFS) and DUD-E-Trans (TransFS) will be discussed here. The results for DenseFS architectures trained on DUD-E-Redocked (RedockedFS) and DUD-E-Hybrid (HybridFS) showed similar trends to TransFS, and are given in Table S3.

### Ligand-Only Test

The precision-recall (PR, left) and receiver-operator characteristic curves (ROC, right) for the Ligand-and-receptor test (green) and Ligand-only test (purple) for the Baseline, OriginalFS or TransFS models are shown in Fig. 1.

**Figure 1:**
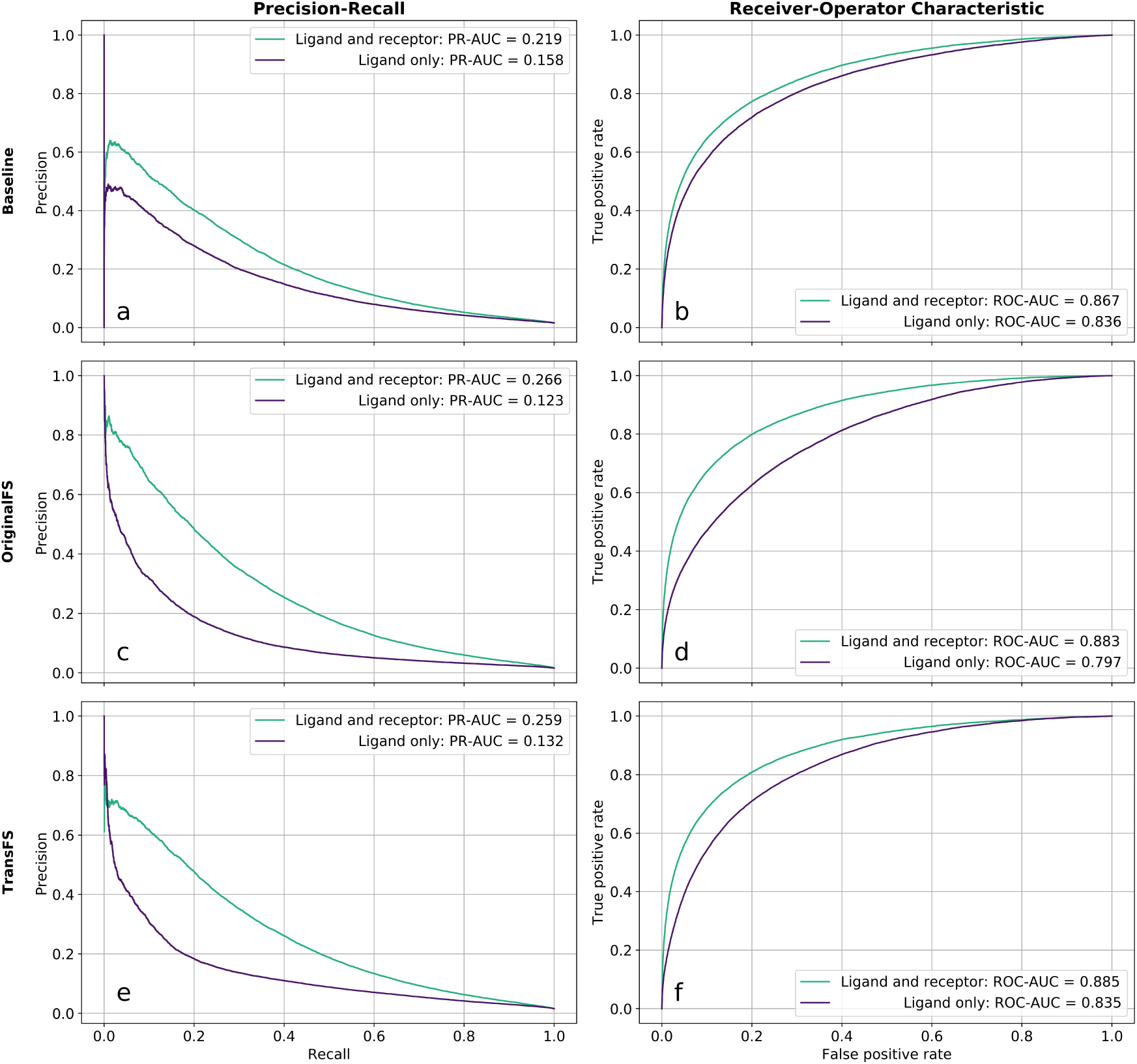
Removing receptor information does not cause total collapse in discriminative power for Gnina or DenseFS-based CNNs. Precision-recall and receiver-operator characteristic curves for the Ligand-and-receptor and Ligand-only tests. Models shown are Baseline (a, b), OriginalFS (c, d) and TransFS (e, f). Green lines indicate held-out DUD-E target test sets with both receptor and ligand information (Ligand-and-receptor test); purple lines are the same set but with only ligand information given (Ligand-only test).

#### Performance of Baseline

Fig. 1 (a, b) shows the PR and ROC curves for Baseline models tested both with and without the receptor present. The difference in the area under the ROC curves (ROC-AUC) between the Ligand-and-receptor and Ligand-only tests is just 0.031 (3.5%, 0.867 to 0.836). This agrees with the previous findings by Chen *et al.*^1^ that performance is hardly affected by the removal of the receptor. The area under the PR curve (PR-AUC) drops by 0.061 (27.9%, 0.219 to 0.158) which is a larger change, but both metrics reflect that a large percentage of predictive ability is still retained in the Ligand-only test. This result indicates that the Baseline model is not relying significantly on information from the receptor channels for its classifications.

#### Performance of OriginalFS

Fig. 1 (c, d) shows the PR and ROC curves for the Ligand-and-receptor and Ligand-only tests, using OriginalFS. The drop in performance when receptor information is removed is far larger than the drop seen for the Baseline. The ROC-AUC drops by 0.086 (9.7%, 0.883 to 0.797) and the PR-AUC decreases by 0.143 (53.8%, 0.266 to 0.123). The larger reduction in predictive ability when the receptor is not present indicates that the more expressive DenseFS CNN architecture uses receptor information more in its classifications than Gnina. However, there is still significant predictive ability in the ligand-only test. OriginalFS is still able to relatively accurately separate actives from decoys using only ligand information.

In an effort to force the DenseFS CNN to learn more from the receptor and how it interacts with the ligand, we trained TransFS using our augmented DUD-E set, DUD-E-Trans. The network therefore sees examples of active ligands in non-physical positions and conformations marked as decoys as well as the correct active poses.

#### Performance of TransFS

Fig. 1 (e, f) shows the PR and ROC curves for the Ligand-and-receptor and Ligand-only tests, using TransFS. The performance is very similar to OriginalFS; the difference in ROC-AUC is 0.050 (5.6%, 0.885 to 0.835) and PRC-AUC drops by 0.127 (49.0%, 0.259 to 0.132). This is surprising, as the TransFS models trained with DUD-E-Trans have seen examples of active ligands in non-physical structures labelled as decoys, which should remove the ability of the network to classify based on ligand fingerprint alone.

For OriginalFs and TransFS there is a significant but similar drop in performance when the receptor is removed. This suggests that using the DUD-E-Trans augmented training set is not causing greater use of the receptor in classification. In order to explore what, if any, influence training using DUD-E-Trans has, we examined the distributions of the raw CNN scores.

The effects of training set on raw CNN scores for actives and decoys in the ligand-only test are shown in Fig. 2. The width of the plots of a particular CNN score is proportional to the relative density of structures assigned that score. Plots are split by training set, whether structures are actives or decoys, and whether a receptor was present (green) or not (purple) in the test set.

**Figure 2:**
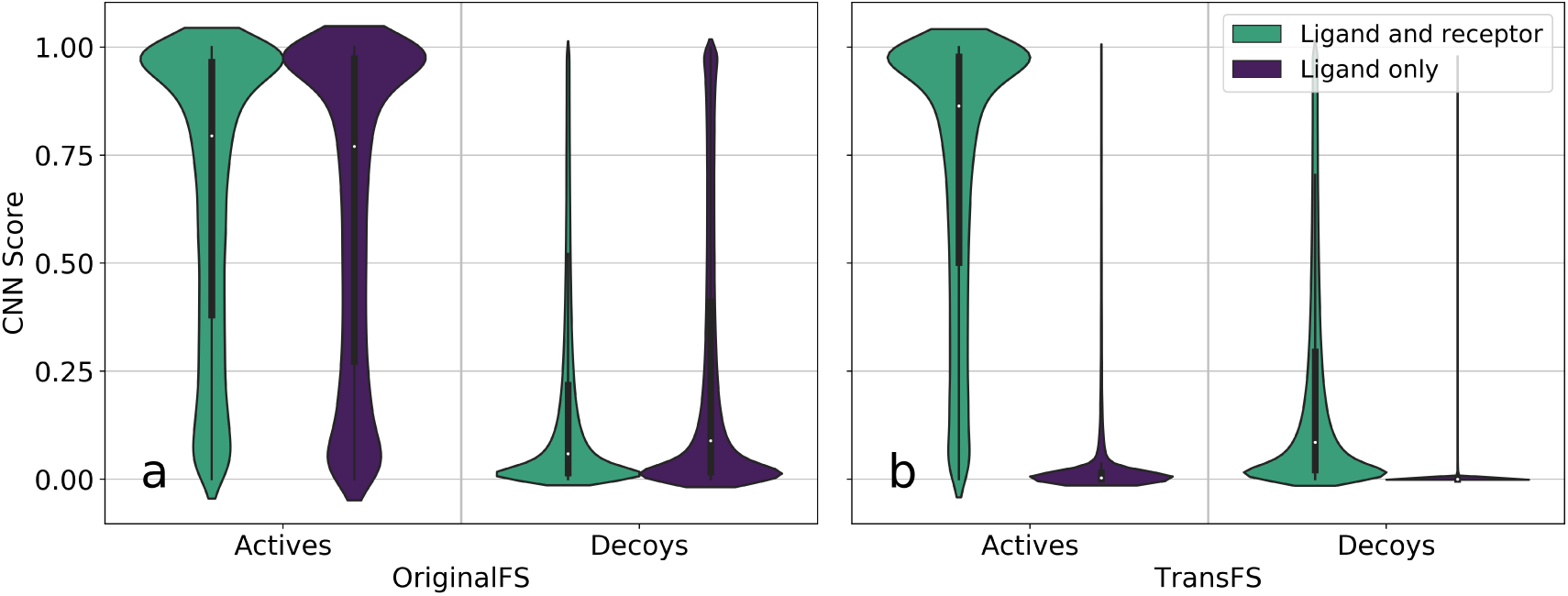
CNN score distributions reveal presence (OriginalFS) and absence (TransFS) of discriminative power when receptor information is removed in Ligand-and-receptor and Ligand-only tests. Violin plots of test set scores given to actives and decoys by OriginalFS and TransFS, with score distributions given with ligand and receptor information present shown in green and with ligand information only in purple. Plots *a* and *b* show distributions for OriginalFS and TransFS CNNs respectively.

#### Score distributions for OriginalFS

Fig. 2a shows the distributions of pose scores given to actives and decoys in the Ligand-and-receptor (green) and Ligand-only (purple) tests by OriginalFS CNNs. Neither actives nor decoys exhibit a significant difference in pose score distribution between the Ligand-and-receptor and Ligand-only tests, i.e. the green (Ligand-and-receptor) and purple (Ligand-only) plots for actives are almost identical. Compared to Baseline (Fig. S4a in the supporting information), decoys in the Ligand-and-receptor test are skewed more towards zero, but actives in the Ligand-only test are in fact skewed more towards higher scores, showing that using the deeper network alone does not guarantee the use of receptor information.

#### Score distributions for TransFS

Fig. 2b shows the same tests as Fig. 2a, using TransFS. Compared with OriginalFS (Fig. 2a), the score distributions for when the receptor and protein (Ligand-and-receptor) are present do not change significantly. However, the distributions for the Ligand-only tests collapse to being highly clustered near zero - for actives, the median score goes from 0.864 to 0.0292 when the receptor is removed, and for decoys the median drops from 0.0855 to 0.000132. These results show that receptor information is being used to generate a significant proportion of the score of the CNN for TransFS.

However, it is not evidence that it is being used in the way that we desire (to pick out physicochemical interactions). In order to test if receptor information is now being used in this way, we next test performance on an external validation set to test for generalisability. As described in the introduction, the ability to better classify structures from a different distribution to the training data would imply that general physical rules have been learned.

### ChEMBL Validation

In order to examine how well the OriginalFS and TransFS CNNs generalize, they were used to predict on the ChEMBL validation set (see methods). The results for each target in the ChEMBL set are given in Table 1.

**Table 1:**
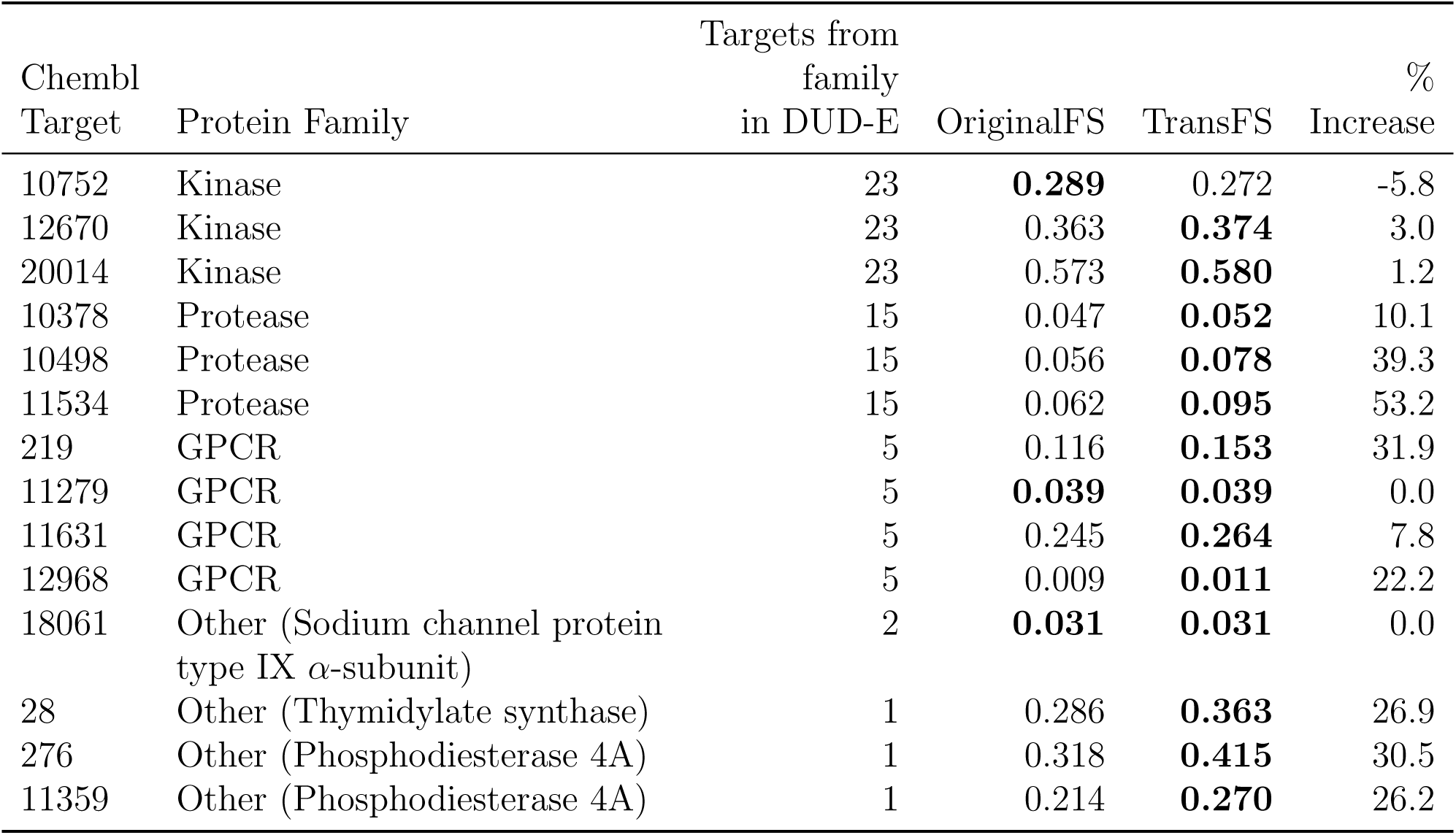
TransFS causes CNNs trained on the DUD-E dataset to generalise better to an external validation set than OriginalFS. PR-AUC values for the 14 targets in the ChEMBL validation set for OriginalFS and TransFS CNNs. Highest values in a row are highlighted in bold.

TransFS performs as well as or better than OriginalFS for 13 out of 14 targets, suggesting that augmentation causes models trained on one dataset to generalize better, and have increased predictive power on a different dataset. For ROC-AUCs, see Table S2 in the supporting information.

Training with DUD-E-Trans (TransFS) rather than DUD-E-Original (OriginalFS) tended to give larger improvements in PR-AUC scores for targets from families which do not feature heavily in the DUD-E set such as proteases (see methods for an overview and Table S1 in the supporting information for a detailed breakdown of DUD-E target families). The largest performance increases were seen for proteins in the ‘Other’ category - containing targets in families such as ‘Phosphodiesterase 4A’ and ‘Thymidylate synthase’ - which do not feature at all in DUD-E. This larger difference in performance for the more distinct targets is to be expected, as ligands which are active for one member of a protein family are likely to be active for others, so learning the fingerprint profile of ligands likely to bind to one member of a family will help performance on external validation sets which contain that family. This can be seen here in the performance on kinase targets, which is extremely close between the two different training regimens. There are 26 kinase targets in the DUD-E training set, and although the ChEMBL set does not contain any identical targets, ligands which are actives for one kinase are much more likely to be active for another. In our case this means that OriginalFS performs comparably on kinases to TransFS, and in the case of the ChEMBL kinase target with ID 10752, OriginalFS has a PRC-AUC that is 0.017 (5.9%) larger than the PRC-AUC achieved by TransFS.

Being better able to generalize to an external validation set, especially on targets from families not in the training set, suggests that more general rules of physical interactions are being learned. In order to look for more direct evidence of this, we used input masking to find which areas of the ligand and protein each CNN assigned importance to when making a classification.

### Masking

Masking can be used to identify which parts of the feature vector are important in the classification given by a CNN, by ‘covering’ parts of the input and comparing the score to the ‘uncovered’ original (see supporting information, Fig. S5). Here, we have used it to check the contribution of each individual atom in both the ligand and receptor. A CNN which learns physical interactions should give high (positive) masking scores to atoms involved favourable interactions such as hydrogen bonding, and negative scores to atoms involved in unfavourable interactions such as steric clashes. Masking was performed as outlined in the methods on PDB structures 10OH and 1W4O, two of the structures used for masking in Hochuli *et al.*^21^ (Fig. 3), neither of which are present in DUD-E.

**Figure 3:**
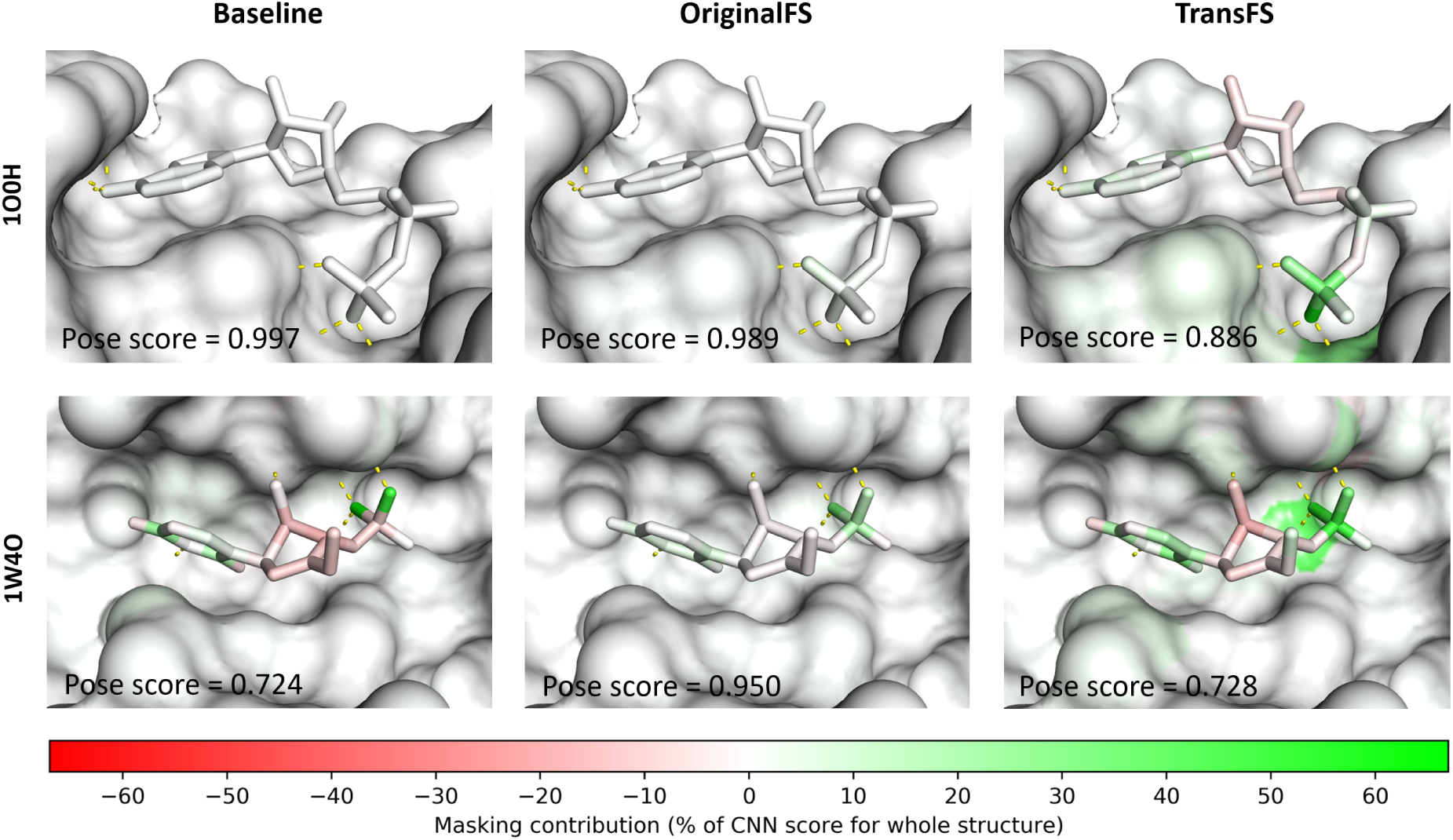
Masking using TransFS CNNs highlights important interactions between the ligand and protein. Results of masking for bound structures with PDB IDs 1O0H (top) and 1W4O (bottom). ned as donor/acceptor pairs within 3.5 °A of one another) are shown as yellow dashed lines. The pose scores from different networks and training sets are shown in the bottom left corners of each image. Color scale shows colors assigned for different masking scores, as a percentage of the pose score of the original unmasked structure.

#### Baseline (left)

There is no contribution displayed by any of the atoms of the receptor for either 1O0H or 1W4O. The highest ligand atom score is 2.4% of the overall CNN score for 1O0H and the two highest scoring atoms for 1W40 are oxygens which appear to be involved in hydrogen bonding to receptor atoms (66.3% and 50.5%).

#### OriginalFS (middle)

A small contribution to the score can be seen for the terminal phosphate of the ligand in 10OH potentially related to hydrogen bonding to the receptor, although this is not matched by coloring in the receptor. The largest contributions for 1O0H are 1.1% (receptor) and 5.6% (ligand). There are larger ligand contributions for 1W4O (max. 34.5%), but again these are not related to interactions sites on the receptor (max. 1.7%).

#### TransFS (right)

There are stronger signals from atoms in the binding sites of the receptors of both structures. The strongest signals from both the ligand and receptor in 1O0H are for atoms involved in hydrogen bonds between the receptor and terminal ligand phosphate group. The largest contributions are 28.6% (receptor, a nitrogen) and 59.8% (a hydrogen-bonding oxygen on the phosphate group in the ligand). The phosphorous atom in terminal group of the ligand is assigned a score of 35.3%. Similar interactions are picked up for 1W4O, again with atoms involved in hydrogen bonds between the ligand and receptor being highlighted as important to the CNN score. Here, the largest contributions are 38.5% (receptor) and 54.9% (ligand).

These results suggest that training on DUD-E-Trans (TransFS) causes CNNs to score receptor and ligand atoms involved in hydrogen bonding, an effect which has not been shown before. They also suggest that networks trained in this way are learning physical interactions, a requirement for a more generalisable classifier.

## Conclusion

In this paper, we have described a method of training data augmentation that improves the generalisability of a deep learning method for structure-based virtual screening. Our augmentation consists of adding three copies of each active in a random conformation, randomly rotated and translated and labelled as decoys for training. Networks trained with this augmented data also have higher scores associated with receptor and ligand atoms that participate in favourable interactions. Our simple augmentation procedure could be configured for use in many other machine learning methods for ligand binding prediction. The arbitrary choice of using three translated decoys for each active could also be further be tuned, for example using cross-validation on a test set external to the training data.

### PDB codes used (PubMed IDs)

1W4O (15670155); 1O0H (14573867)

