## Supporting information for "Dataset Augmentation Allows Deep Learning-Based Virtual Screening To Better Generalize To Unseen Target Classes, And Highlight Important Binding Interactions"

E-mail:

Protocol S1: Protocol for generation of DUD-E-Trans augmented training set.

First the active ligand molecules are 'moved' into the centre of mass of the protein, then a distance is generated by the following method ($R$ = radius of the protein):

$$\begin{aligned}
y_1(r) &= \exp\left(-\frac{(r-\mu_1)^2}{2\sigma_1^2}\right)\\
y_2(r) &= \exp\left(-\frac{(r-\mu_2)^2}{2\sigma_2^2}\right)\\
y_3(r) &= \exp\left(-\frac{(r+\mu_2)^2}{2\sigma_2^2}\right)\\
y_4(r) &= 2y_1(r) + y_2(r) + y_3(r)\\
\text{where} \quad \mu_1 &= 0 \quad \sigma_1 = \frac{R}{3.5}\\
\mu_2 &= R \quad \sigma_2 = \frac{R}{2.5}
\end{aligned}$$

$y_4(r)$ is calculated for a range of 1,000 values of $r$ in the interval $(-1.3R, 1.3R)$. The set of values for $y_4(r)$ is then divided by the total sum of those values, making a discrete distribution over $r$ from which the distance to move the molecule in the direction of a random unit vector is drawn. The OpenEye toolkit[S1] is used to randomly rotate and then randomize the conformation of each translated molecule. These new active/target structures are labelled as decoys for training.

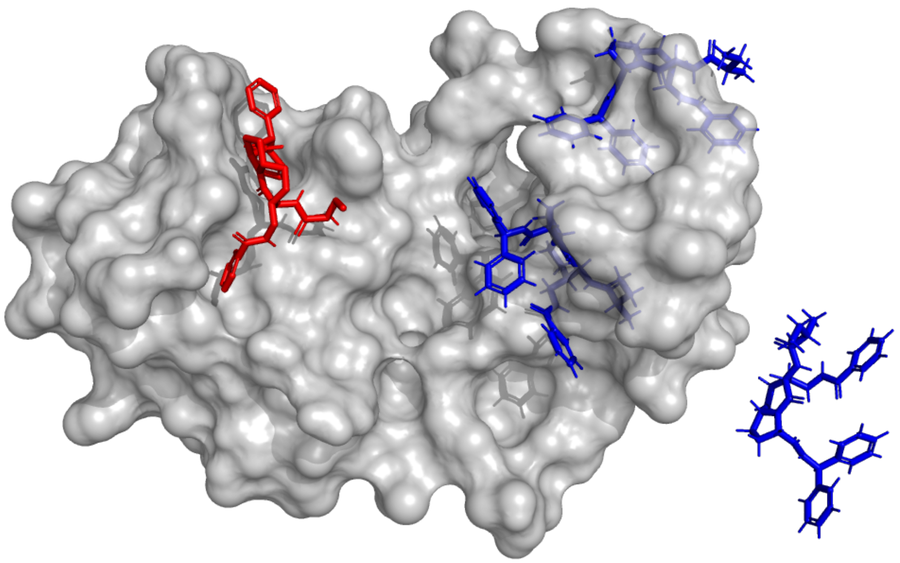

Figure S1: Example of random translations and conformations (shown in blue) of an active molecule (original pose in red) for a protein target (grey), included in the DUD-E-Trans dataset. Blue poses are labelled as decoys in a DUD-E-Trans training set, with red remaining as an active. DUD-E target: XIAP; ligand: CHEMBL584393.

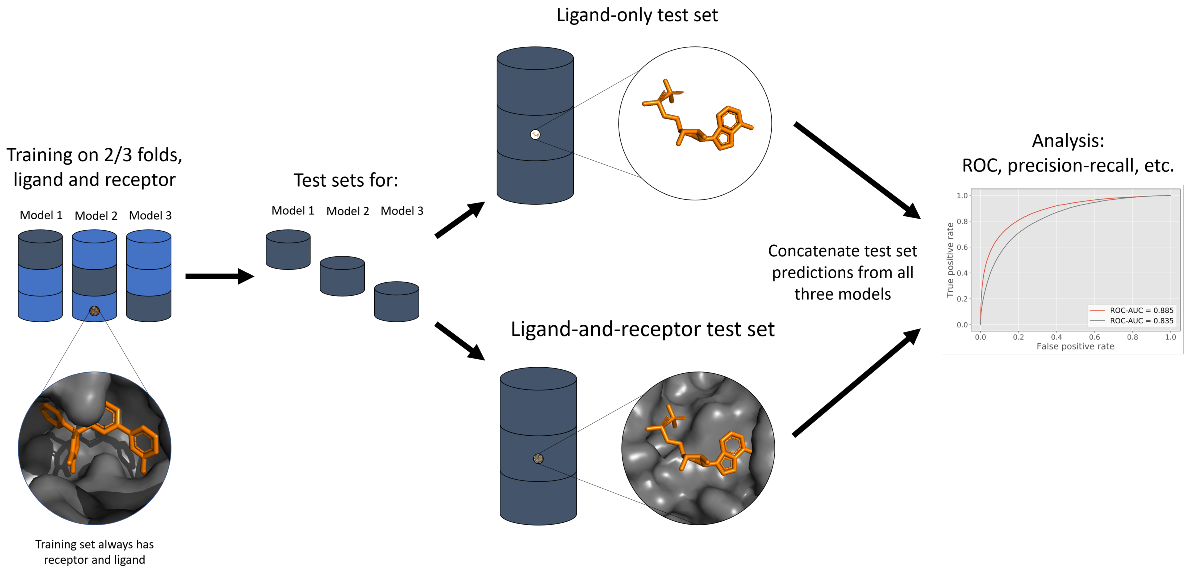


Figure S2: Protocol for training on DUD-E and the Ligand-and-receptor and Ligand-only tests. Three models are trained on 2 folds each (light blue), and tested on both normal and ligand-only versions of the final fold (dark blue). Test predictions from the three models are concatenated for analysis.

Table S1: Target splits for the DUD-E dataset. Targets from different folds have $\leq 80\%$ sequence similarity.

| Fold 0 | | Fold 1 | | Fold 2 | |
|---|---|---|---|---|---|
| Target | Family | Target | Family | Target | Family |
| abl1 | Kinase | akt1 | Kinase | csf1r | Kinase |
| braf | Kinase | akt2 | Kinase | kit | Kinase |
| cdk2 | Kinase | egfr | Kinase | mapk2 | Kinase |
| fak1 | Kinase | jak2 | Kinase | mk14 | Kinase |
| igf1r | Kinase | lck | Kinase | mp2k1 | Kinase |
| kpcb | Kinase | met | Kinase | plk1 | Kinase |
| mk01 | Kinase | tgfr1 | Kinase | rock1 | Kinase |
| mk10 | Kinase | wee1 | Kinase | vgfr2 | Kinase |
| src | Kinase | ace | Protease | mmp13 | Protease |
| ada17 | Protease | bace1 | Protease | lkha4 | Protease |
| casp3 | Protease | fa7 | Protease | reni | Protease |
| dpp4 | Protease | hivpr | Protease | try1 | Protease |
| tryb1 | Protease | fa10 | Protease | gcr | Nuclear |
| urok | Protease | thrb | Protease | mcr | Nuclear |
| andr | Nuclear | esr1 | Nuclear | ppara | Nuclear |
| ppard | Nuclear | esr2 | Nuclear | prgr | Nuclear |
| drd3 | GPCR | pparg | Nuclear | rxra | Nuclear |
| tysy | Other | thb | Nuclear | aa2ar | GPCR |
| hdac8 | Other | cxcr4 | GPCR | adrb1 | GPCR |
| hivrt | Other | aces | Other | adrb2 | GPCR |
| pur2 | Other | pyrd | Other | glcm | Other |
| aofb | Other | pgh1 | Other | cah2 | Other |
| inha | Other | parp1 | Other | grik1 | Other |
| comt | Other | cp2c9 | Other | ital | Other |
| sahh | Other | def | Other | dhi1 | Other |
| pygm | Other | pnph | Other | fpps | Other |
| fabp4 | Other | pgh2 | Other | pde5a | Other |
| aldr | Other | ada | Other | nos1 | Other |
| fnta | Other | cp3a4 | Other | kif11 | Other |
| pa2ga | Other | nram | Other | hivint | Other |
| xiap | Other | fkb1a | Other | hxk4 | Other |
| hmdh | Other | ptn1 | Other | kith | Other |
| dyr | Other | hdac2 | Other | ampc | Other |
| | | gria2 | Other | hs90a | Other |

| | Model 1 | Model 2 | Model 3 | Pose mean |
|---|---|---|---|---|
| **Docked pose 1** | 0.93 | 0.91 | 0.95 | 0.93 |
| **Docked pose 2** | 0.99 | 0.90 | 0.87 | 0.92 |
| **Docked pose 3** | 0.92 | 0.94 | 0.90 | 0.92 |
| **Docked pose 4** | 0.83 | 0.85 | 0.78 | 0.82 |
| **Docked pose 5** | 0.84 | 0.71 | 0.88 | 0.81 |
| **Docked pose 6** | 0.78 | 0.77 | 0.76 | 0.77 |
| **Docked pose 7** | 0.74 | 0.72 | 0.64 | 0.70 |
| **Docked pose 8** | 0.66 | 0.66 | 0.66 | 0.66 |
| **Docked pose 9** | 0.54 | 0.60 | 0.68 | 0.61 |

| Ligand score |
|---|
| 0.88 |

Figure S3: How a final CNN score is arrived upon given three scores for nine different poses of the same ligand for each target. Each pose is scored by three different models trained in the same way from a different starting configuration; the mean of these three scores is taken, giving nine scores (one per pose), the top five of which are averaged to give the final score.

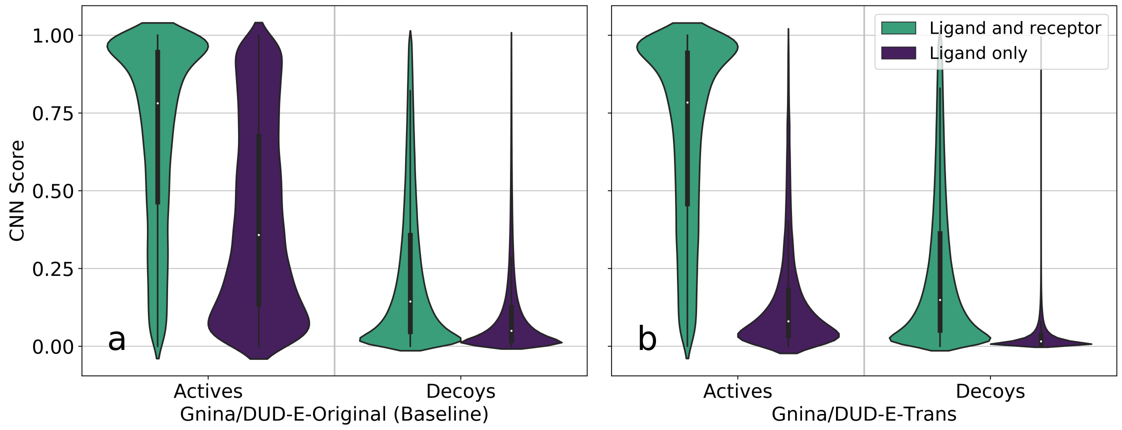


Figure S4: Violin plots of scores given to actives and decoys by Gnina models trained with DUD-E-Original (Baseline) and DUD-E-Trans. Score distributions for test sets with ligand and receptor information present are shown in green; test sets with ligand information only are shown in purple. Plots *a* and *b* show distributions for models trained using DUD-E-Original and DUD-E-Trans respectively.

Table S2: ROC-AUC values for the 14 targets in the ChEMBL validation set for DenseFS models trained either with DUD-E-Original or with the DUD-E-Trans dataset. Highest values in a row are highlighted in bold.

| Chembl Target | Family | OriginalFS | TransFS |
|---|---|---|---|
| 10752 | Kinase | **0.875** | 0.854 |
| 12670 | Kinase | **0.912** | 0.910 |
| 20014 | Kinase | 0.934 | **0.938** |
| 10378 | Protease | 0.636 | **0.705** |
| 10498 | Protease | 0.743 | **0.769** |
| 11534 | Protease | 0.804 | **0.842** |
| 219 | GPCR | 0.842 | **0.865** |
| 11279 | GPCR | 0.819 | **0.825** |
| 11631 | GPCR | **0.845** | 0.828 |
| 12968 | GPCR | 0.505 | **0.582** |
| 28 | Other (Thymidylate synthase) | **0.927** | 0.914 |
| 276 | Other (Phosphodiesterase 4A) | 0.868 | **0.882** |
| 11359 | Other (Phosphodiesterase 4A) | 0.834 | **0.840** |
| 18061 | Other (Sodium channel protein type IX $\alpha$-subunit) | 0.677 | **0.693** |

Table S3: PR-AUC values for the 14 targets in the ChEMBL validation set for DenseFS models trained with the DUD-E-Original, DUD-E-Trans, DUD-E-Redocked and DUD-E-Hybrid datasets.

| Chembl Target | Family | OriginalFS | TransFS | RedockedFS | HybridFS |
|---|---|---|---|---|---|
| 10752 | Kinase | 0.289 | 0.272 | 0.270 | 0.272 |
| 12670 | Kinase | 0.363 | 0.374 | 0.350 | 0.335 |
| 20014 | Kinase | 0.573 | 0.580 | 0.566 | 0.525 |
| 10378 | Protease | 0.047 | 0.052 | 0.032 | 0.041 |
| 10498 | Protease | 0.056 | 0.078 | 0.073 | 0.091 |
| 11534 | Protease | 0.062 | 0.095 | 0.100 | 0.103 |
| 219 | GPCR | 0.116 | 0.153 | 0.069 | 0.087 |
| 11279 | GPCR | 0.039 | 0.039 | 0.040 | 0.044 |
| 11631 | GPCR | 0.245 | 0.264 | 0.191 | 0.202 |
| 12968 | GPCR | 0.009 | 0.011 | 0.011 | 0.011 |
| 28 | Other (Thymidylate synthase) | 0.286 | 0.363 | 0.297 | 0.323 |
| 276 | Other (Phosphodiesterase 4A) | 0.318 | 0.415 | 0.269 | 0.375 |
| 11359 | Other (Phosphodiesterase 4A) | 0.214 | 0.270 | 0.133 | 0.195 |
| 18061 | Other (Sodium channel protein type IX $\alpha$-subunit) | 0.031 | 0.031 | 0.032 | 0.033 |

Table S4: PRC-AUC values for the 14 targets in the ChEMBL validation set for OriginalFS and DenseFS trained on the entire of their respective training sets, both with and without receptor information available at test time. The right-most column shows results for the TrasnFS models given only ligand information to predict on, and scores are significantly lower than for the corresponding test with OriginalFS.

| ChEMBL Target | Family | OriginalFS Ligand and receptor | OriginalFS Ligand only | TransFS Ligand and receptor | TransFS Ligand only |
|---|---|---|---|---|---|
| 10752 | Kinase | 0.289 | 0.300 | 0.272 | 0.069 |
| 12670 | Kinase | 0.363 | 0.348 | 0.374 | 0.060 |
| 20014 | Kinase | 0.573 | 0.548 | 0.580 | 0.261 |
| 10378 | Protease | 0.047 | 0.059 | 0.052 | 0.089 |
| 10498 | Protease | 0.056 | 0.059 | 0.078 | 0.076 |
| 11534 | Protease | 0.062 | 0.062 | 0.095 | 0.058 |
| 219 | GPCR | 0.116 | 0.078 | 0.153 | 0.123 |
| 11279 | GPCR | 0.039 | 0.032 | 0.039 | 0.016 |
| 11631 | GPCR | 0.245 | 0.213 | 0.264 | 0.095 |
| 12968 | GPCR | 0.009 | 0.009 | 0.011 | 0.009 |
| 28 | Other | 0.286 | 0.295 | 0.363 | 0.193 |
| 276 | Other | 0.318 | 0.272 | 0.415 | 0.054 |
| 11359 | Other | 0.214 | 0.162 | 0.270 | 0.032 |
| 18061 | Other | 0.031 | 0.029 | 0.031 | 0.012 |

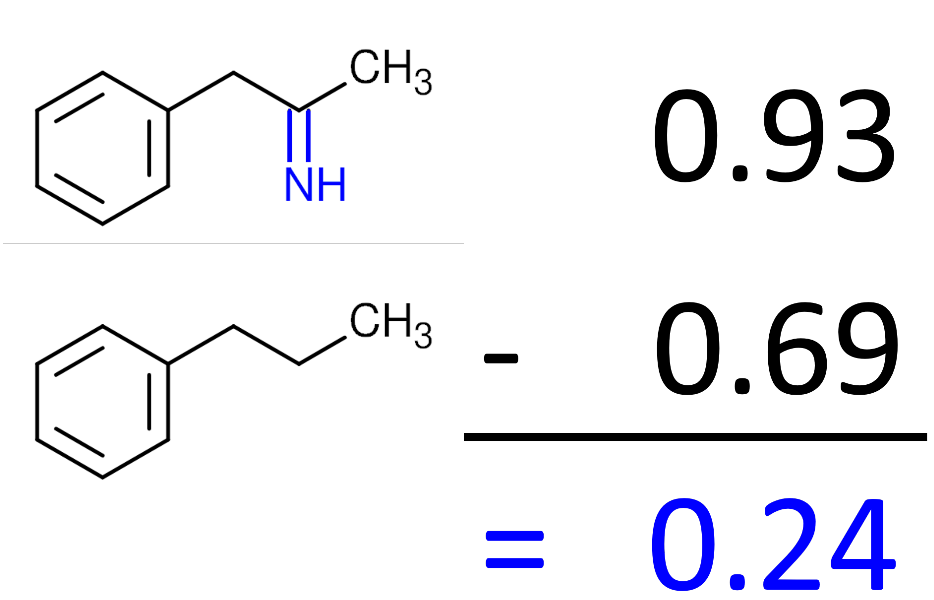


Figure S5: Atomistic masking. The contribution of each atom is given by the difference in scores when the atom is present compared to when it is not.
